# APV9WR: An Integrated Web Resource of Alpha papillomavirus 9 for Genomics, Proteomics, Phylogenetic and Therapeutic Analysis

**DOI:** 10.1101/2024.06.28.601125

**Authors:** Akanksha Kulshreshtha, Vasu Goel, Akriti Verma, Sparsh Goel, Susha Dinesh, Sameer Sharma, Ratul Bhowmik, Ashok Aspatwar

**Author notes:** **Corresponding author** Dr. Ashok Aspatwar.

## Abstract

**Purpose:** Alphapapillomavirus 9 is a virus belonging to the Papillomaviridae family. It has a close genetic relationship with high-risk HPV-16 and other HPV strains such as HPV-31, HPV-52, HPV-35, HPV-58, HPV-67, and HPV-35. This virus is responsible for causing warts and malignant tumors and is responsible for about 75% of cervical malignancies and pre-cancerous lesions worldwide. As a result, it requires specialized research and attention. Our goal is to create a comprehensive resource that can assist researchers and scientific groups in their work.

**Material and Methodology:** A total of 1230 full genome sequences and 9140 protein sequences were obtained from GenBank and NCBI Virus, respectively. Further Phylogenetic Analysis, Codon usage and context analysis, CpG islands analysis, Glycosylation sites, Diagnostic Primers, B cell Epitopes and MHC I and MHC II binders were identified and analyzed using relevant Bioinformatics tools and Python program

**Results:** APV9WR is a web resource that was developed after analysis. Our data indicate that HPV 35 and HPV 38 have the most genomic diversity. From codon usage analysis, it was observed that AAA, AUU, UAU, UGU, and UUU are the most used codons, while ACG, CCG, CGA, CGG, CGU, GCG, and UCG are some of the unusual codons in APV9 nucleotide sequence with accession id - LC626346.1. We found 4714 CpG island locations in 1230 complete nucleotide sequences of APV9, and only 663 CpG island locations were unique. Further N-linked glycosylation, O-linked glycosylation, diagnostic primers, Potential B-cell epitopes and MHC I and MHC II binders were also analyzed and tabulated.

**Conclusion:** We have consolidated basic information about the virus, such as entire genomic sequences and proteins. It primarily comprises a wide range of studies and outcomes, including genome alignment, phylogenetic inferences, codon context and usage bias, and important CpG island statistics. Furthermore, primers for molecular diagnostics were identified, and glycosylation sites were located and investigated. Most significantly, potential therapeutic elements such as vaccine epitopes and obtaining potential information about them were investigated. Our collective effort on this tool is meant to serve the greater good of the research community for therapeutic intervention for Alphapapillomavirus 9. Using this link https://apv9nsut.web.app will take you to the web app.

## 1. INTRODUCTION

Human papillomaviruses (HPVs) are small, non-enveloped, double-stranded DNA viruses that preferentially infect epithelial cells of the vaginal and upper respiratory tracts, and skin. There are approximately 150 types of HPV that have been identified and classified into different genera based on their DNA sequences [1]. HPV types have been classified into five major genera: alpha (65 different types), beta (54 types), gamma (99 types), mu (03), and nu (1type) [2]. Epidemiological evidence designated alphapapillomaviruses (alpha HPVs) as high-risk HPV types [3]. Alphapapillomavirus 9 (APV9) is one of 14 APV species. Its virion is non-enveloped, icosahedral, tiny, and 60 nm in size [4]. APV9 has three genomic regions. The Long Control Region contains an early promoter and regulatory element that is crucial to viral DNA transcription and replication; the Early Region contains proteins (E1, E2, E4, E5, E6, and E7) involved in viral gene expression, replication, and survival; and the Late Region contains proteins that are crucial to viral replication, survival, and gene expression [3]. APV9 represents several subtypes namely HPV-16, HPV-33, HPV-58, HPV-52, HPV-31, HPV-35, and HPV-67 [5]. Genital warts, papilloma, and cancer etc. are associated with these species. Around 75% of cervical cancer cases in women are caused by APV9 [6]. E6 and E7 are oncoproteins that modifies host immune response pathways to generate a chronic infection in cervical cancer. They also change cells. The scientific community prioritizes research into this virus due to its high carcinogenicity, although no stable vaccination is available till now. Gardasil 9, a recombinant non-covalent vaccination, protects against nine HPV types: 6, 11, 16, 18, 31, 33, 45, 52, and 58. Current HPV vaccines have safety, uncertainty, and cost-effectiveness issues are there. HPV research has been ongoing for a long time, and efforts have been made to create a resource for HPV therapeutics [7], but no complete resource exists for APV9, the species responsible for most cervical cancer cases, to aid researchers and the scientific community. Realizing the significance and need to have more understanding of this virus, an integrated resource was created to help the research community decipher APV9 as shown in figure 1. Further, this resource covers various kinds of analysis and results which were derived using bioinformatics software or conceptually written codes. Genome alignment, phylogenetic inferences, codon context and usage bias, CpG islands, and glycosylation sites are available to researchers. B-cell vaccine epitopes and possible MHC-I and MHC-II binders have also yielded promising findings.

**Fig. 1:**
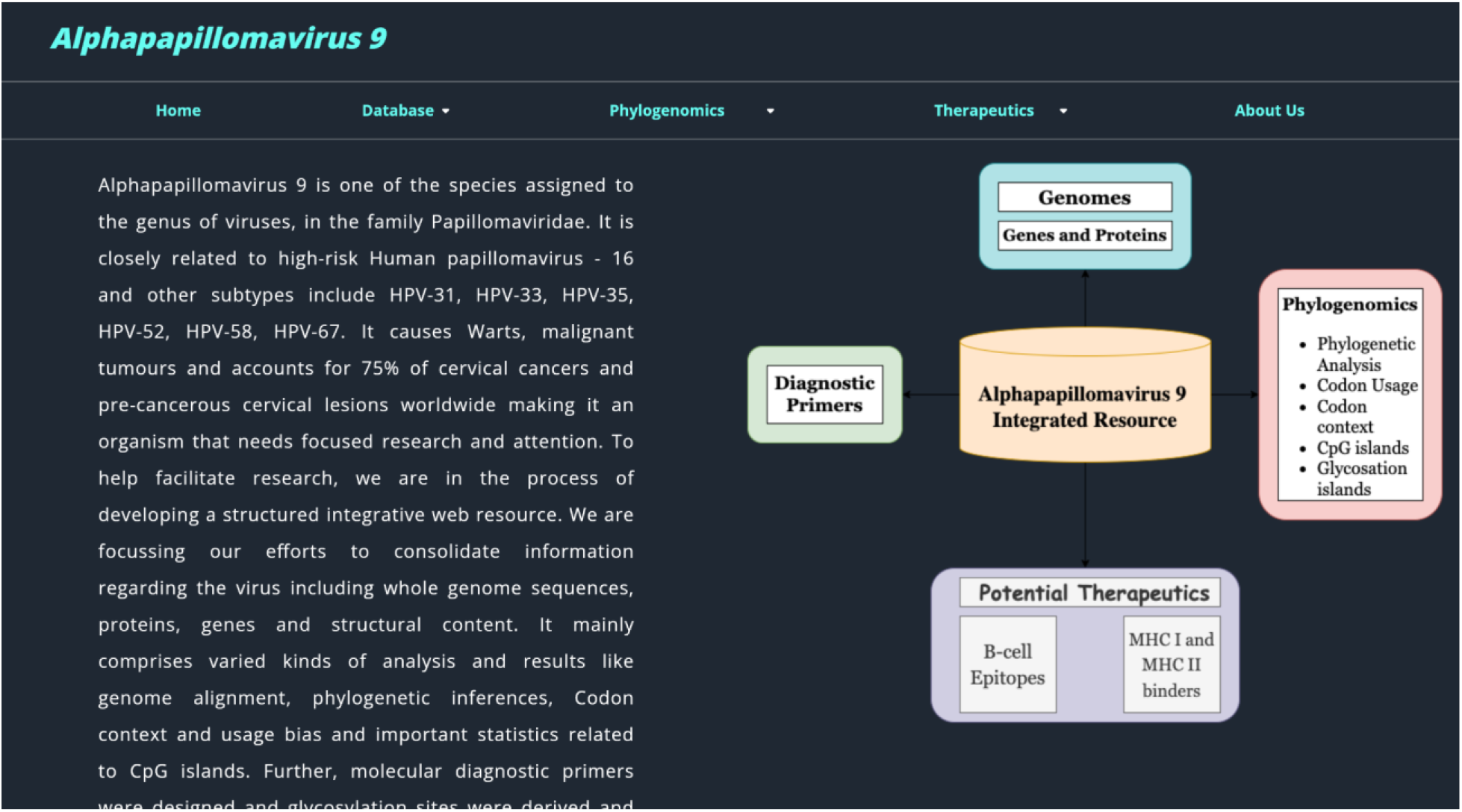
Overview of the resource

## 2. MATERIALS AND METHODOLOGY

### 2.1 Genomic & Proteomic Data Collection and Clustering

APV9’s genomic and proteomic sequences were taken from GenBank and NCBI Virus for study. 1230 full genome sequences and 9140 protein sequences were obtained till October 2021 and segregated according to continents. The following information was collected about the organism: strain, isolation, genome size, geographical area, host, region, and year, as shown in the Supplementary file, Table 1. To reduce the number of protein sequences to be evaluated, the CD-HIT Suite grouped them [8]. Clustering was based on an identity percentage greater than or equal to 99%, which resulted in 583 unique clusters of protein sequences (Supplementary file, Table 2).

**Table 1:** Few predicted B cell epitopes with their score generated by LB tope software (SVM score), percentage probability for a correct prediction, respective nucleotide accession no. and protein name.

| S.No | Accession No | Protein Name | Epitope | SVM Score | B cell confidence |
| --- | --- | --- | --- | --- | --- |
| 1 | BCU08869.1 | E1 | IVKLLRYQNVEFTQF | 1.1717 | 89.06 % |
| 2 | BCU08870.1 | E2 | KEKQAPAKRRRPDVP | 0.8828 | 79.43 % |
| 3 | BCU08871.1 | E4 | APKKNRRLPNDDDLT | 1.1868 | 89.56 % |
| 4 | BCU08872.1 | E5 | FCIVLRPLLLSIYVY | 0.7279 | 74.26 % |
| 5 | BCU08867.1 | E6 | MFQDTDEKPRNLHEL | 0.8322 | 77.74 % |
| 6 | BCU08868.1 | E7 | MDRPDGQAKPETTNY | 0.9065 | 80.22 % |
| 7 | BCU08874.1 | L1 | GCKPPTGEHWGKGTP | 1.4207 | 97.36 % |
| 8 | BCU08873.1 | L2 | MDPDTPFPQPSISYS | 0.9641 | 82.14 % |

**Table 2:**
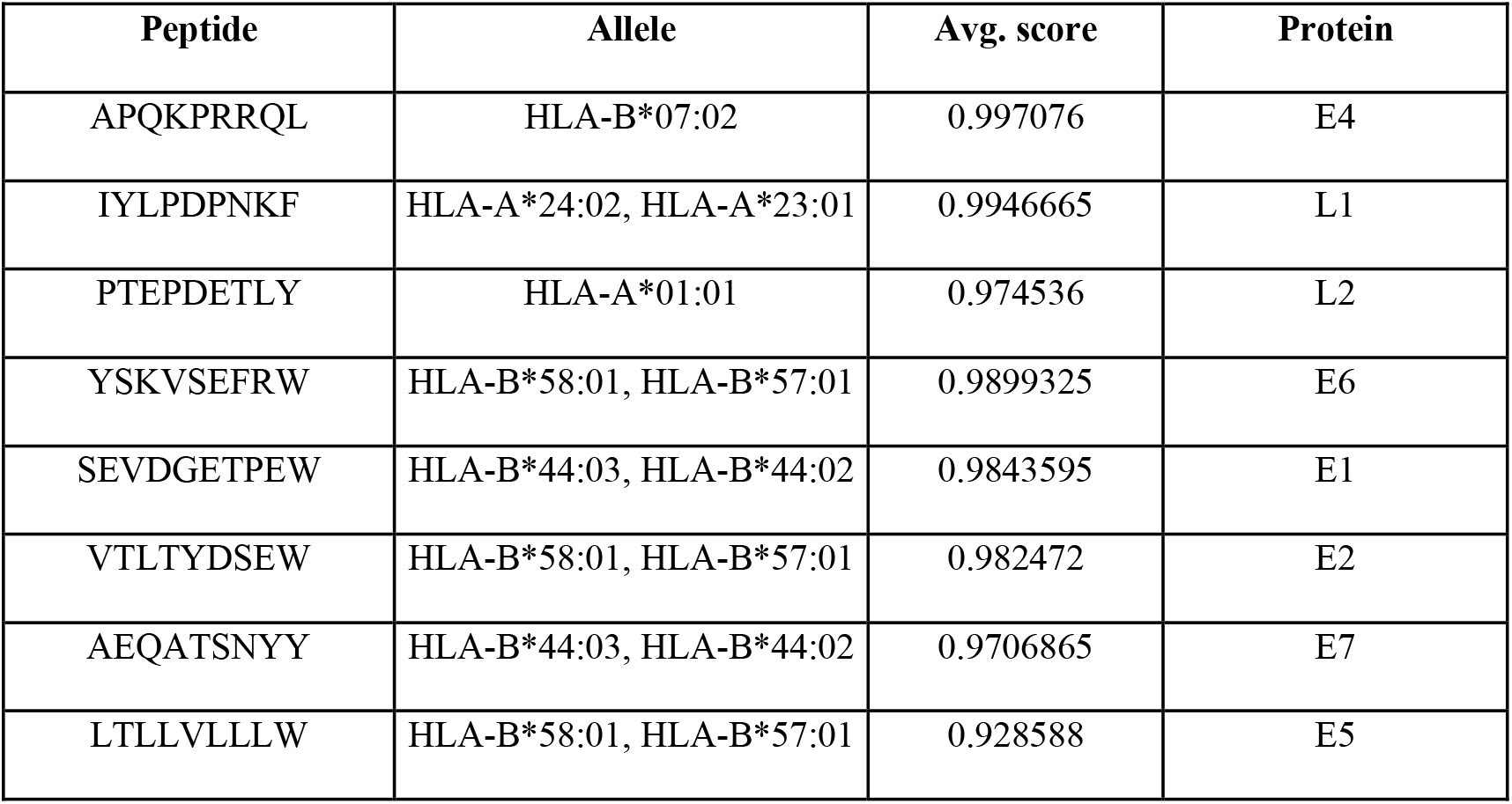
Representation of protein-wise predicted MHC I binders along with their average scores.

### 2.2 Phylogenetic Analysis

APV9’s evolution and HPV subtype relationships were examined using a phylogenetic tree. Based on an identity percentage greater than *99%*, CD-HIT Suite [8] produced 90 groups from 1230 genome sequence isolates from the NCBI database. Multiple Sequence alignment was done on each representative genome of 90 clusters using MAFFT online, too [9]. Phylogenetic trees were generated using MEGA 11 [10]. The p-distance substitution model created neighbor-joining trees. Bootstrapping with 100 pseudo-replicates increased dependability. Another phylogenetic tree was created using the inbuilt feature at the NCBI Virus site [11], which matched HPV subtype evolution to their country.

### 2.3 Codon usage and context analysis

Codon bias is measured by row frequency (the frequency by which a codon is used for each amino acid). Anaconda program [12] was used to look at the distribution of rare and preferred codons as well as codon context among APV9 strains.

### 2.4 CpG islands analysis

Gardiner-Garden and Frommer’s [13] approach for identifying and analyzing CpG island locations was used. A nucleotide window of at least 200 bp traveling at 1 base pair interval throughout the sequence is used to calculate. CpG islands are sequence ranges with an observed/expected value of CpG sites more than 0.60 and GC content greater than 50%. The estimated value of CpG dimers in a window is calculated by dividing the product of the window’s Gs and Cs by its length. CpG islands are often found in vertebrate genes’ 5’ regions. Therefore, this methodology can also identify genomic sequences’ putative genes.

### 2.5 Glycosylation

NetCGlyc v1.0 [14] was used to identify C-linked glycosylation sites in APV9 proteins envelope (1,2,4,5,6,7), Early Protein, L- (1&2), Major Capsid, Minor Capsid, Regulatory, Replication, and Transforming. NetOGlyc v4.0 [15] and NetNGlyc-1.0 [16] use neural networks to identify O- and N-linked glycosylation sites respectively.

### 2.6 Diagnostic Primers

For finding out the primers of the sequences, NCBI’s tool named Primer-Blast [17] at default conditions was used; it is a combination of two software, primer3, and blast, which enables the user to design and check specific primers.

### 2.7 B cell Epitopes

For APV9, LBtope software [18] was used to predict linear B-cell epitopes to help in finding potential therapeutic targets. LBtope uses Support Vector Machine based models. It is an efficient method with an accuracy of ∼81% built on a large experimentally verified database of epitopes and non-epitopes.

### 2.8 MHC I and MHC II binders

MHC class I and class II binders were anticipated for vaccine design in this investigation. IEDB Analysis Resource Consensus [19] was used for predictions. IEDB’s consensus technique (CombLIb, SMM, and ANN) assessed MHC I binders. IEDB guidelines define high binding affinity as IC values < 50 nM. Results with a score 0.5 and a low percentile rank indicate high binding affinity. The IEDB consensus technique (SMM-align, NN-align, Sturniolo, and CombLib) predicted MHC-II binders. For high-affinity binders in the output table, the rank cutoff was 0.4.

## 3. RESULTS AND DISCUSSION

### 3.1 Phylogenetic Analysis

The Neighbor-joining method’s ensuing phylogenetic tree (Fig. 2) enables us to comprehend type-specific variations within the genomes analyzed. Our data indicate that HPV 35 and HPV 38 have the most genomic diversity. HPV 58 and HPV 33 were closely connected. To summarize, two major groups, including similar subtypes, are Group A (HPV 35, HPV 31, HPV 16, Alpha-9) and Group B (HPV 58, HPV 33, HPV 67, HPV 52). Further, another phylogenetic tree was constructed to determine the link between isolates from various nations. 96 sequences with legitimate isolation sources were taken for this study, and the resulting phylogenetic tree (Fig. S1) demonstrated that nations share comparable subtypes of species due to their geographic proximity. Additionally, the relationship between subtypes and countries was analyzed: HPV 35 is prevalent in Africa, South America, and a few Asian nations (India, Thailand, and Georgia), HPV 16 is prevalent in Brazil, India, and Nepal, while HPV 52 and HPV 33 are prevalent in Brazil.

**Fig. 2:**
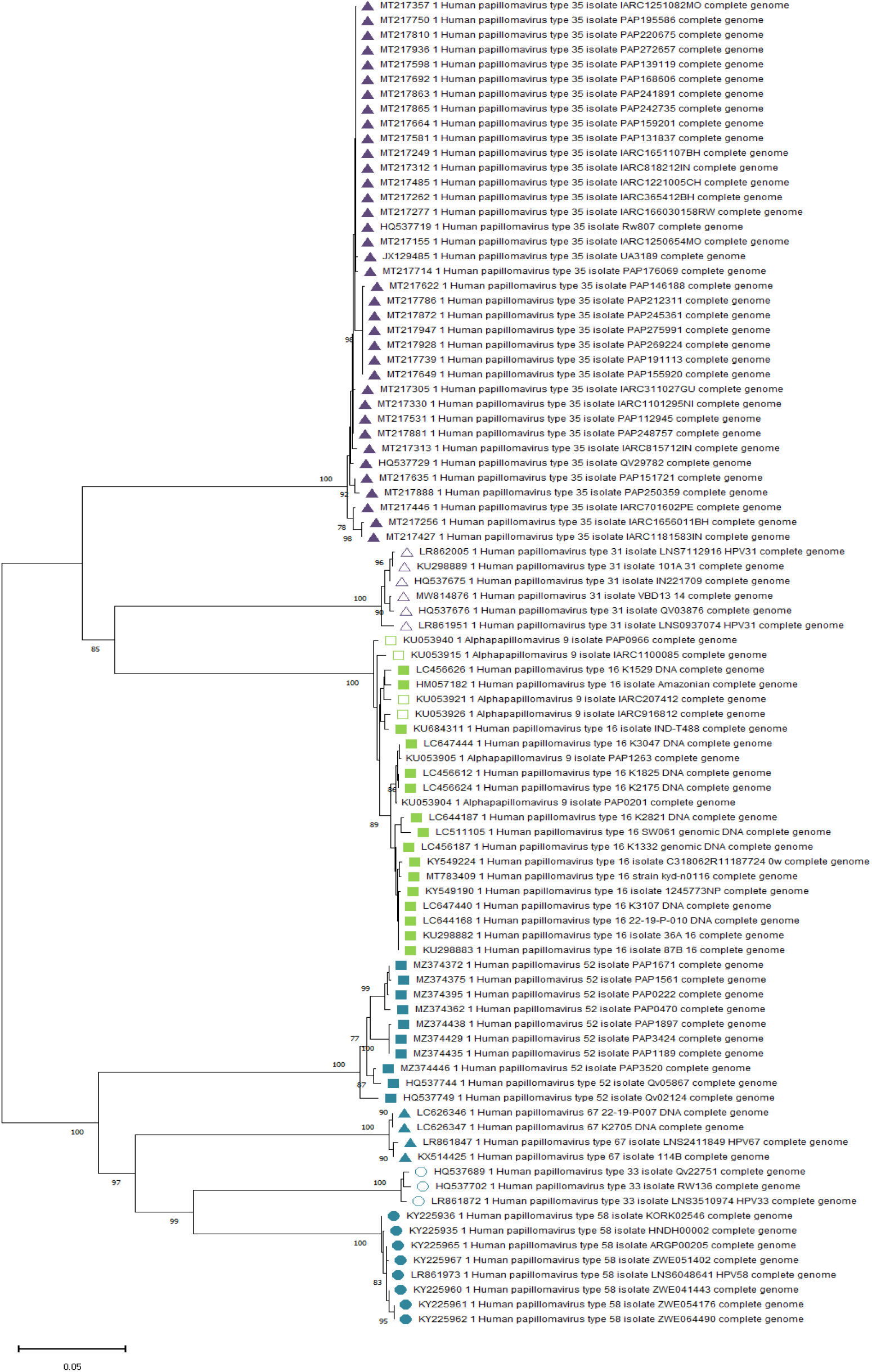
The evolutionary history was inferred using the Neighbor-Joining method conducted in MEGA11. The percentage of replicate trees in which the associated taxa clustered together in the bootstrap test (100 replicates) are shown next to the branches for values > 75%. The midpoint rooted tree is drawn to a scale 0.05. The tree was computed using the p-distance method and is in the units of the number of base differences per site. Codon positions included were 1st+2nd+3rd+Noncoding.

### 3.2 Codon usage and context analysis

Codon usage varies between genomes. It’s affected by nucleotide composition bias, gene length, recombination events, expression level, and G+C content. This uneven use of synonymous codons may help uncover species-specific genome evolution patterns. anaconda program calculates genome sequence residual values for each codon pair. Average residual values for all codon pairs were calculated. A two-colored heat map (fig. 3) was created for each frequency table cell value. In the heat map, red and green indicate rare and favored codons. The heat-cluster map’s pattern shows codon context differences and similarities between species. The cell’s black color indicates non-biases-related codon context residual values. Blue indicates infrequent codons, and black indicates favored codons in a histogram. The histogram in Supplementary File Fig. 2 was created using the APV9 nucleotide sequence with accession number LC626346.1. AAA, AUU, UAU, UGU, and UUU are the most regularly used codons, while ACG, CCG, CGA, CGG, CGU, GCG, and UCG are rare codons.

**Fig. 3:**
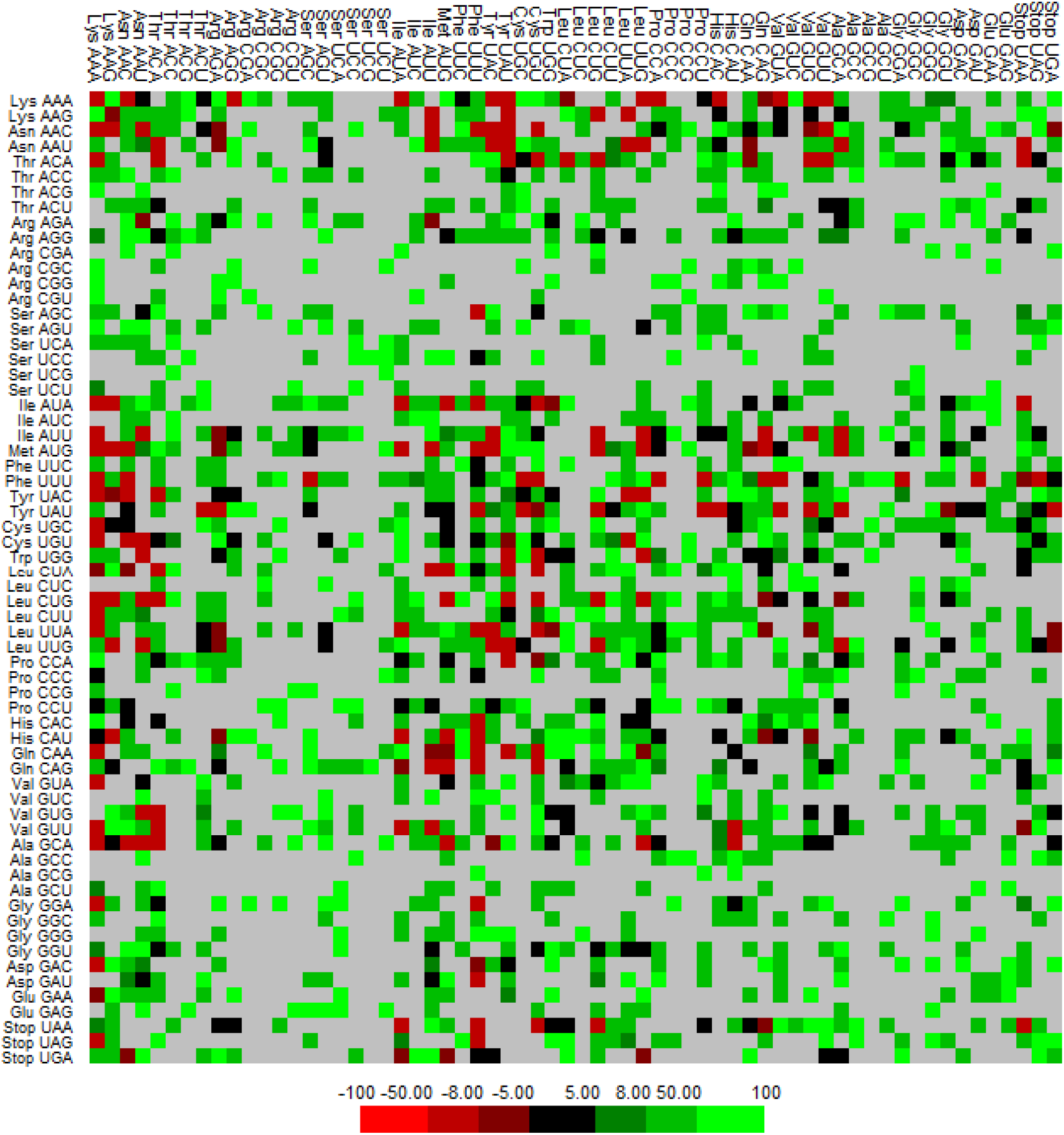
Two-colored heat map generated for each value in a frequency table cell. Green and red colors represent preferred and rare codons, respectively, in the matrix. The black color of the cells denotes residual values related to codon context that does not match biases.

### 3.3 CpG islands analysis

CpG islands (CGIs) have several CpG dinucleotide repeats (CpG sites). Guanine and cytosine nucleotides follow each other in the linear sequence of CpG sites. CpG islands are sequence regions with at least 200 bp, GC content larger than 50%, and Obs/Exp greater than 0.6. The estimated value of CpG dimers in any window is calculated by dividing the product of the window’s Gs and Cs by its length. A python script found CpG island sites in the provided nucleotide by moving a 200-bp window at 1-bp intervals throughout the sequence. Python program results are in Table 3 of the supplementary file. 4714 CpG islands were discovered in 1230 APV9 nucleotide sequences. We evaluated these 4714 CpG island positions and discovered that only 663 were unique, the longest was 378 bp, the average GC content was 50%, and the maximum Obs/Exp ratio was 0.90.

### 3.4 Glycosylation

All NCBI-listed APV9 strains were analyzed for O-, N-, and C-linked glycosylation sites. Viruses use N-linked glycosylation for protein trafficking, receptor binding, immunological evasion, virus assembly, pathogenicity, and more [20]. N-linked glycosylation is most prevalent in proteins E1(31,53,272,314), E2(21,83,84,133,134,219,252), E6 at 80, E7(5,29,71), L1 at 216, and L2 (143,149,159,160,208,209,394,397,419). O-linked glycosylation sites are involved in signal transduction, ligand recognition, virus/bacteria-host interactions, and more [21]. The most prevalent O-linked glycosylation sites in proteins are E1(168,167,166), E2(207,247,264), E4 at 47, E6 at 6, and L2 at (133,134,156,212,213). C-linked glycosylation sites may affect enzyme activity and secretion [22]. We found no protein C-linked glycosylation sites were not found in any protein as 0.5 0.5 thresholds of the software (For reference and results).

### 3.5 Diagnostic Primers

To analyze the diagnostic primers of, 1230 genome sequence isolates obtained from the NCBI database,23 clusters were formed using CD-Hit Suite [8] based on identity percentage greater or equal to 90%. Each genome sequence from the cluster data was run on Primer-Blast software to get primers for each sequence. This was repeated for 23 sequences, and then common primers from the data generated were taken out. Supplementary file, Table 4 shows all the primer sequences.

### 3.6 Potential B-cell epitopes

The current study predicts and consolidates plausible B-cell epitopes to create vaccination candidates after integrating MHC binders. 9140 protein sequences from the NCBI viral database were grouped with 99% identity into 583 distinct clusters. We detected 2459 non-redundant B-cell epitopes from 583 entries. Our website (https://apv9nsut.web.app) lists 2459 epitopes with accession number, protein name, predicted epitope, Lbtope SVM score, and B-cell confidence. Table 1 lists a few epitopes that may be targets for each protein to help explain the results.

### 3.7 MHC I and MHC II binders

3429 MHC I and 903 MHC II binders were predicted from 583 representative protein sequences. These recommended data, including projected vaccine epitopes, can aid vaccine design. For each projected binder, the website (https://apv9nsut.web.app) provides epitope sequence, start and finish position, score, and percentile rank. The screenshots of integrated web resource are depicted in supplementary file (fig.3-fig.9)

## 4. Conclusion and Future Prospects

APV9-associated carcinogenicity is a topic of concern as it is the main cause of 75% of cases of invasive cervical cancer recorded worldwide, and a systematic and comprehensive approach may pave the way for the rapid design and development of a successful strategy against carcinogenic APV9. There are currently just a few in silico studies on APV9, and no such database or resource exists. The integrated web resource of APVP resource is an effort toward building a platform for the researchers that covers various components of phylogenomics like phylogenetic analysis, codon usage, codon context, CpG islands, and glycosylation sites. Our findings reflect a significant advancement in the APV9 studies, providing an analysis of potential B-cell epitopes and MHC I and MHC II binders, which can assist in vaccine designing and development. Further, the predicted molecular primers listed in our resource will help discover the potential diagnostic techniques.

## Abbreviations

APV9: Alphapapillomavirus 9
MHC: Major Histocompatibility Complex
HPV: Human papillomaviruses

## Acknowledgments

**The authors are very much thankful to the authority of NSUT, New Delhi**, for moral support and encouragement.

## Statements and Declaration

The authors have no conflicts of interest to declare. All co-authors have seen and agree with the contents of the manuscript, and there is no financial interest to report. We certify that the submission is original work and is not under review at any other publication.

## Author contribution statement

Conception and planning: VG, AV, SG, RB, and AK; Data collection and processing: VG, AV and SG: Analysis performed: VG, AV, and SG; Prepared figures/graphs and table; VG, AV, and SG; Database implementation; VG Manuscript drafting, editing and revision; VG, AV, RB, AA, SS and SG; Supervision and funding acquisition AA: All the approved final version of the manuscript.

## Ethics approval

The research does not include any clinical samples or human samples; hence, Ethical approval is not applicable, and no clinical trials were done.

## Data Availability

- \**Supplementary files for further information and* for further assistance refer to the website https://apv9nsut.web.app.

